# lncRNA-screen: an interactive platform for computationally screening long non-coding RNAs in large genomics datasets

**DOI:** 10.1101/087080

**Authors:** Yixiao Gong, Hsuan-Ting Huang, Yu Liang, Thomas Trimarchi, Iannis Aifantis, Aristotelis Tsirigos

**Author notes:** To whom correspondence should be addressed. Aristotelis Tsirigos (AT). Correspondence may also be addressed to Iannis Aifantis (IA). Present Address: Aristotelis Tsirigos, Department of Pathology, New York University School of Medicine, New York, New York, 10016, USA.

## Abstract

Long non-coding RNAs (lncRNAs) have emerged as a class of factors that are important for regulating development and cancer. Computational prediction of lncRNAs from ultra-deep RNA sequencing has been successful in identifying candidate lncRNAs. However, the complexity of handling and integrating different types of genomics data poses significant challenges to experimental laboratories that lack extensive genomics expertise. To address this issue, we have developed lncRNA-screen, a comprehensive pipeline for computationally screening putative lncRNA transcripts over large multimodal datasets. The main objective of this work is to facilitate the computational discovery of lncRNA candidates to be further examined by functional experiments. lncRNA-screen provides a fully automated easy-to-run pipeline which performs data download, RNA-seq alignment, assembly, quality assessment, transcript filtration, novel lncRNA identification, coding potential estimation, expression level quantification, histone mark enrichment profile integration, differential expression analysis, annotation with other type of segmented data (CNVs, SNPs, Hi-C, etc.) and visualization. Importantly, lncRNA-screen generates an interactive report summarizing all interesting lncRNA features including genome browser snapshots and lncRNA-mRNA interactions based on Hi-C data. In summary, our pipeline provides a comprehensive solution for lncRNA discovery and an intuitive interactive report for identifying promising lncRNA candidates. lncRNA-screen is available as free open-source software on GitHub.

## INTRODUCTION

The landscape of transcription in organisms is now known to be complex and pervasive, producing a wide range of small and long RNA species with a variety of biological functions discovered so far (1–3). One of the least characterized yet largest class of RNA species are the long noncoding RNAs (lncRNAs). The most basic definition of a lncRNA is a long RNA, at least 200 base pairs in length, that does not encode protein. They can be further classified by features such as their genomic location, structure, and expression (4). Of the small number of lncRNAs that have been characterized, they have been shown to be functionally important for chromatin and other cellular processes that affect organismal development and cancer (5, 6). However, there are 28,031 lncRNA transcripts annotated to date on GENCODEv19 and 91,000 in MiTranscriptome, and this number will continue to grow as deeper and more sensitive RNA sequencing data are generated. Recently, large-scale lncRNA analyses of published data (e.g. TCGA) have been conducted (7, 8), however the authors have not made their pipelines available. A number of databases and bioinformatics tools have been developed to annotate and catalog lncRNAs that are known or novel (9). LncRNA2Function (10), LNCipedia (11), lncRNAdb (12) and lncRNAtor (13) provide comprehensive databases for known, annotated lncRNAs. iSeeRNA (14), CPC (15) and CPAT (16) introduced machine learning-based approaches only focusing on assessment of the coding probability of potential lncRNAs. lncRScan (17) and its new version lncRScan-SVM (18) are pipelines provide novel multi-exonic lncRNA only discovery from RNA-seq, lacking the ability to integrate other data types to further filter for interesting lncRNA candidates. It is known that lncRNAs, like coding genes, are enriched for histones that mark transcriptionally active sites such as histone H3K4me3 and H3K27ac (19, 20). This approach has led to the successful identification of LUNAR1 and its function in regulating the IGF1R locus to sustain T-cell leukemia growth (21). Thus integrating other genomic features allows for identification of lncRNAs with important biological functions. One of the fundamental issues in the field is to identify the function of the discovered lncRNAs. In this context, the ability to integrate a variety of genomic datasets to increase the probability of identifying lncRNAs that are functionally relevant will be of great value. Thus having an extensive bioinformatics pipeline that can quickly annotate and classify lncRNAs from RNA sequencing (RNA-seq) data will be valuable for identifying strong candidates for biological validation. To this end, we have developed an extensive computational pipeline to integrate different types of experimental data to annotate lncRNAs, which can be filtered by the user to identify specific lncRNAs of interest. The pipeline first takes RNA-seq data to align and assemble to build a comprehensive transcriptome assembly for all the samples. Then, using a series of filtering criteria based on gene annotations, sequence length, expression level, coding potential and other features, a list of putative lncRNA candidates is defined containing basic information that includes transcript size, genomic location, and differential gene expression. Afterwards, depending on the raw data that users may provide, our pipeline can process and annotate the putative lncRNAs with other processed information such as ChIP-seq data, copy number variation, Hi-C interaction etc. Gene tracks for UCSC genome browser and lncRNA local genomic snapshots are generated simultaneously in order to quickly visualize and assess each lncRNA by its features. *The output of the pipeline is a comprehensive table and an interactive HTML report containing all the putative lncRNAs with their corresponding genomic features* that the user has added into the pipeline. This report can then be filtered interactively by the user for specific lncRNAs of interest based on any combination of the genomic features included in the analysis.

## MATERIAL AND METHODS

### The lncRNA-screen workflow

lncRNA-screen is an extensive analysis pipeline providing various useful functions for lncRNA annotation and candidate selection. It encompasses multiple automated processes designed for lncRNA discovery and computational selection, including public data download, locally sequenced data integration (RNA-seq and ChIP-seq datasets), comprehensive transcriptome assembly, coding potential estimation, expression level quantification, differential expression comparisons and analysis, and lncRNA classification. Our pipeline enables fully customizable lncRNA discovery with insightful visualization embedded in each step of data processing. Most importantly, lncRNA-screen automatically generates an interactive lncRNA feature report that allows users to conveniently search, filter, and rank by important features (e.g. expression level, presence of histone marks, etc.) extracted from the different input data types. Additionally, it provides a genome snapshot of each lncRNA locus to help users visually assess the relevance and quality of each candidate lncRNA. The main functionality of the lncRNA-screen pipeline compared to other published lncRNA analysis tools is summarized in Table 1. According to the table, lncRNA-screen provides the most extensive computational lncRNA discovery pipeline to date. The lncRNA-screen workflow can be divided into two main phases (Figure 1). In Phase I, the pipeline first conducts RNA-seq alignment and assembly individually for each sample and then merges them in order to construct a comprehensive transcriptome assembly. Next, a series of filtration steps are applied in order to detect lncRNAs based on reference annotations and genomic location. A putative lncRNA candidate list is generated in Phase I for further classification in Phase II using different data types. In Phase II, expression levels of the putative lncRNAs are quantified in all the samples and summarized at multiple user-defined group levels. Moreover, by integrating with ChIP-seq data, we identify typical histone mark profiles (H3K4me3, H3K27ac) around the TSS region of all lncRNAs. Pairwise differential expression analysis can also be performed between groups. At the same time, lncRNA-screen annotates all putative lncRNAs using the information obtained from the analysis and supplemented by additional types of “segmented” data (CNVs, SNPs, Hi-C, etc.) in order to generate a comprehensive interactive lncRNA feature report. Users can easily adjust parameters (see *README.md* file on GitHub repository) to find the most confident and functionally relevant lncRNAs candidates for further experimental validation. Furthermore, lncRNA-screen automatically produces different summary plots and a flowchart providing intuitive guidance to the users for adjusting the parameters. Finally, for every lncRNA, lncRNA-screen generates a genome snapshot which shows the gene structure, expression pattern and histone mark profile for the neighboring area. Additionally, if processed Hi-C data is provided, a local Hi-C interaction snapshot revealing potential looping events between each lncRNA and its neighboring genes is also generated.

**Table 1.** The function comparison between lncRNA-screen and other public available software.

|  | lncRNA-screen | coRAL | RNA-CODE | lncRScan | iSeeRNA | CIRI | Annocript | LncRNA2Function |
| --- | --- | --- | --- | --- | --- | --- | --- | --- |
| Stand-alone server | √ | √ | √ | √ | √ | √ | √ |  |
| **Parallel computing** | √ |  |  |  |  |  |  |  |
| Online server |  |  |  |  | √ |  |  | √ |
| Raw data support | √ |  | √ | √ | √ | √ | √ | √ |
| Processed data support | √ |  |  |  |  | √ |  |  |
| **TCGA, GEO, SRA data automatic download** | √ |  |  |  |  |  |  |  |
| Known lncRNA identification | √ | √ | √ |  |  |  |  |  |
| Novel lncRNA identification | √ | √ |  | √ | √ |  |  |  |
| Differential expression analysis | √ | √ | √ | √ | √ | √ |  |  |
| **ChIP-seq histone mark integration** | √ |  |  |  |  |  |  |  |
| **Hi-C data integration** | √ |  |  |  |  |  |  |  |
| **lncRNA genome snapshots** | √ |  |  |  |  |  |  |  |
| **Interactive lncRNA report** | √ |  |  |  |  |  |  |  |

**Figure 1.**
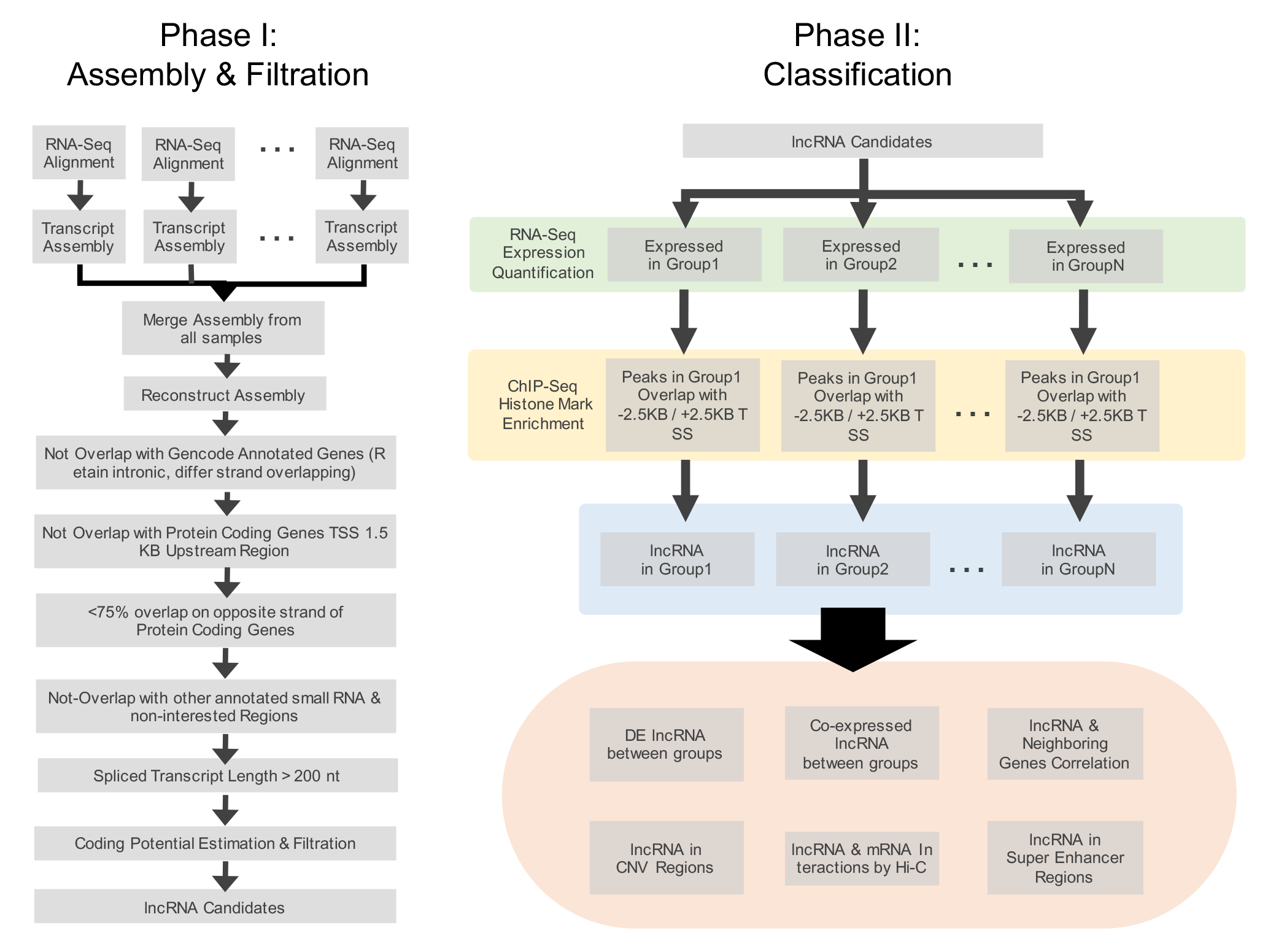
The workflow of lncRNA-screen. Phase I conducts RNA-seq alignment and transcriptome assembly, performs transcript filtration and generates a putative lncRNA list. Phase II uses RNA-seq expression quantification, ChIP-seq histone marks and user-defined annotations (CNVs, Hi-C, etc.) to classify lncRNA into different groups.

### Input data, sample sheet and group information setup

lncRNA-screen provides fully automated and parallel download of raw RNA-seq fastq files from different public data repositories including the Sequence Read Archive (SRA) from the National Center for Biotechnology (NCBI) and The Cancer Genome Atlas (TCGA). The user only needs to provide a list of SRA accession numbers or TCGA UUIDs matched with user-defined sample names to be used throughout the analysis in a sample sheet file. The pipeline automatically downloads the relevant files, while the various tools within our pipeline automatically identify them as the appropriate inputs for downstream analyses. Processed ChIP-seq histone mark data (H3K4me3, H3K27ac and H3K4me1) is required in BED4 format where the fourth column corresponds to the ChIP enrichment score, calculated by MACS2 or any other peak calling tool. Any type of data that can be represented as segmented data in BED4 format (e.g. CNV or SNP data) is also supported as input to lncRNA-screen. For example, copy-number variation segments can be integrated by lncRNA-screen to identify lncRNAs located in recurrently amplified or deleted regions in cancer samples. A group information sheet is also required, where rows correspond to sample names and different grouping strategies can be designated by adding an arbitrary number of columns. Matched histone mark data for each group are also assigned using this file. lncRNA-screen will automatically perform the analysis for all user-defined grouping strategies.

### Read quality assessment for RNA-seq and ChIP-seq

FastQC is a commonly used package for assessing the Next-generation sequencing read quality. lncRNA-screen utilizes FastQC as an automatic quality control (QC) procedure for each sample. It reports the distribution of average per-base and per-sequence quality, per-base and per-sequence GC content, sequence length distribution, sequence duplication level, etc. This information allows users to quickly diagnose irregularities in their input samples and take appropriate action. Based on the results, users can decide whether to perform pre-processing of the fastq files such as trimming during the next step.

### Read alignment

Next, all sequences are aligned using the Spliced Transcripts Alignment to a Reference (STAR) software (22). A pre-built STAR genome index is required in advance. For each sample, the pipeline identifies all the raw reads belonging to each sample, automatically determines whether they are single-end or paired-end and properly groups read pairs as well as different sequence files originating from different sequencing lanes. Then, STAR will automatically determine the strand specificity and read length. This is a significant advantage compared to Tophat2 (23) and other aligners, given that it is oftentimes a challenging task to retrieve this information from large public datasets such as TCGA or SRA. Determining this automatically using STAR is also useful in subsequent steps, for example during transcriptome assembly performed by Cufflinks. Trimming, soft clipping and other type of necessary pre-preprocessing of the reads can be done by STAR during the alignment process. Only uniquely mapped reads are retained in the final alignment. Raw read counts are generated at the same time by the STAR aligner for all the annotated features in the GENCODEv19 GTF as default annotation file (or other annotation files user supplied as a parameter) provided when aligning the samples.

### Post-alignment assessment and processing

After alignment, Picard-Tools (http://broadinstitute.github.io/picard) are used to assess the duplication rate and remove duplicate reads if necessary. We then generate read coverage signal track files in BIGWIG format with adjustable resolution compatible with IGV and UCSC genome browsers. The pipeline also provides a function which can merge and generate combined track files at the group level. An interactive HTML RNA-seq analysis report is generated automatically which incorporates an alignment quality report (example shown in Figure 2; see Results for details) allowing the users to quickly inspect the alignment rates and the number of usable reads. A sample distance plot (example shown in Figure 3; see Results for details) represents an unbiased clustering of the samples based on genes annotated by GENCODEv19. If the number of groups provided by the users is sufficiently small (up to 10), the pipeline also performs differential expression analysis for each pair of groups and reports all the GENCODEv19 annotated genes that pass a user-defined significance threshold. This interactive RNA-seq report is designed for providing an overview of the sample differences and similarities between and within groups and for verifying that the user-defined list of genes follow the expected expression pattern across different groups.

**Figure 2.**
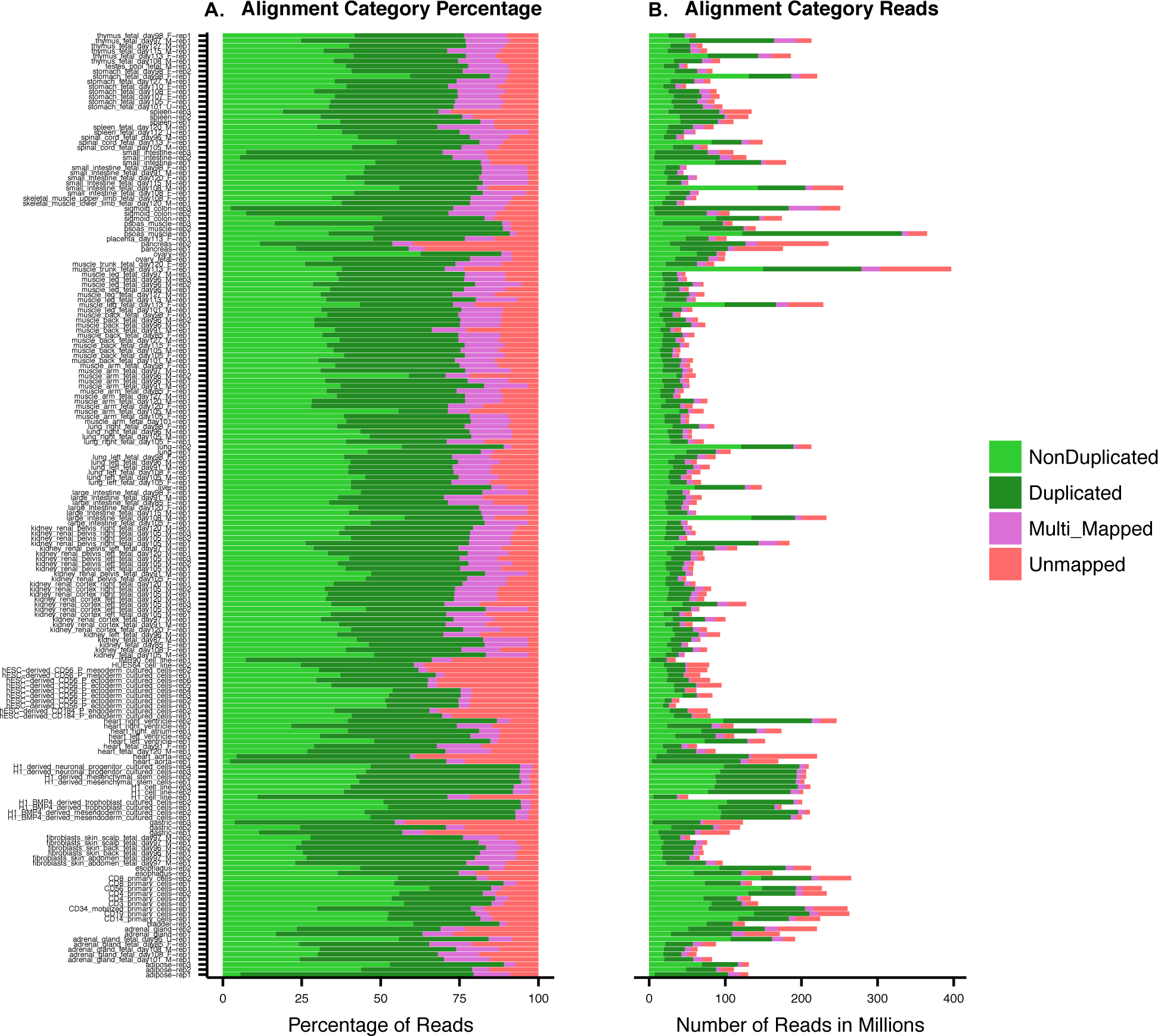
RNA-seq alignment quality plot: (A) read percentages, (B) read counts.

**Figure 3.**
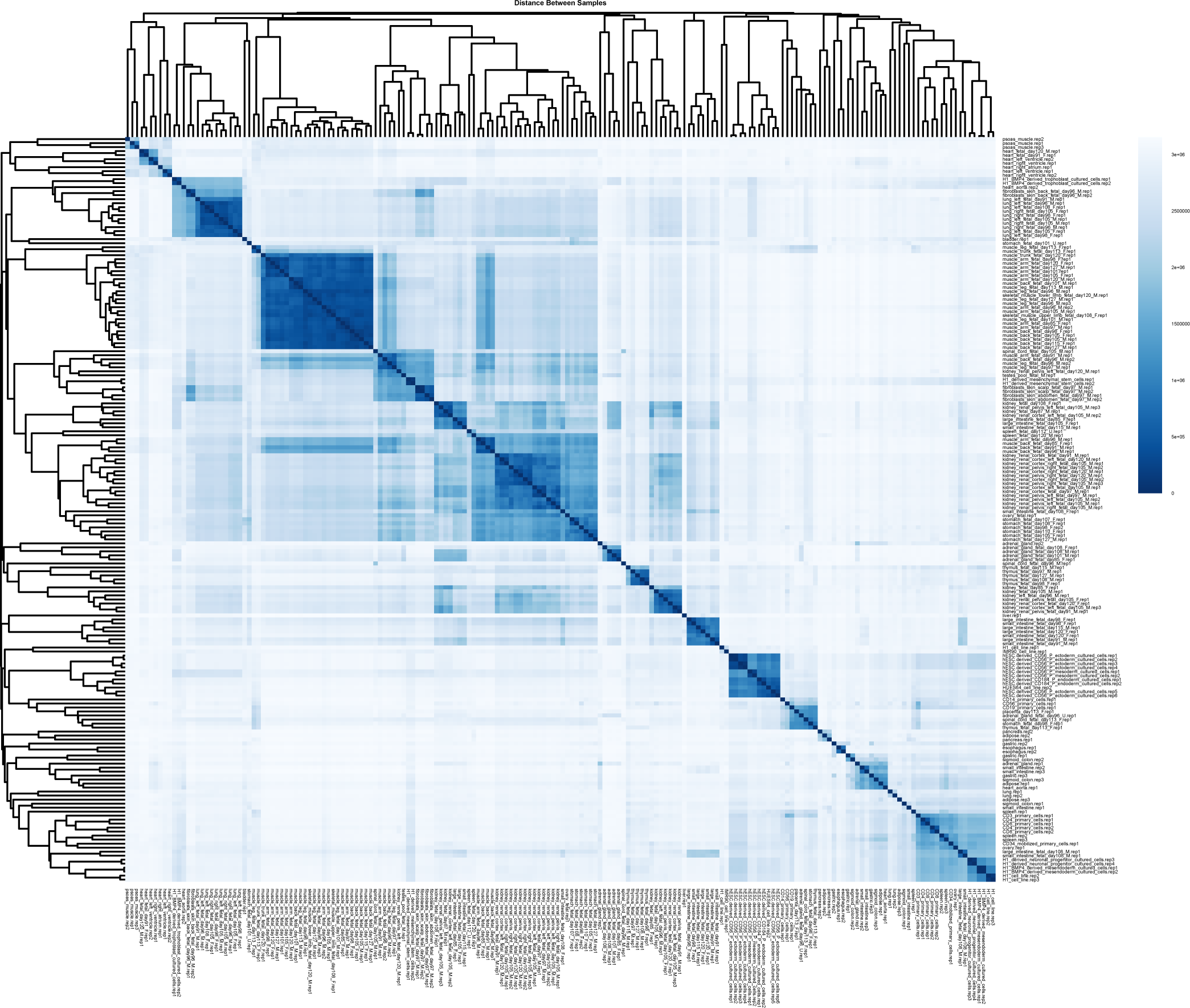
Sample distance report shows unbiased clustering of the samples based on the ENSEMBL annotated genes.

### Transcriptome assembly construction

Aligned sequences are assembled individually by Cufflinks 2.2.1 (24) using NCBI RefSeq annotation (25, 26) as a guide transcriptome with default parameters. All ribosomal RNAs and snRNAs are masked. The strand specificity is automatically determined by the pipeline based on the result of STAR aligner. Finally, we use Cuffmerge to merge all the assemblies into a single comprehensive assembly from all assembled transcripts with GENCODEv19 annotation as a guided assembly.

### Comprehensive identification of putative lncRNAs

To distinguish known transcripts from novel transcripts, we use the Cuffcompare result generated during Cuffmerge process, which compares the comprehensive assembly from all assembled samples and GENCODEv19 reference annotation. Based on the Cuffcompare classification result, all of the transcripts are categorized into 12 different classes. By default, user can define which categories to keep. By default, we keep the three following categories: a transfrag falling entirely within a reference intron (class code “i”); unknown, intergenic transcript (class code “u”); An intron of the transfrag overlaps a reference intron on the opposite strand; exonic overlap with reference on the opposite strand (class code “x”). Annotated lncRNAs by GENCODEv19 will be added back to the filtered transcripts, forming a comprehensive annotated and novel lncRNA assembly. Then this lncRNA assembly is merged at gene level based on the unique gene ID given by Cuffmerge. Alternatively, spliced transcripts for each gene can be retrieved in the future if necessary by searching the entire transcript annotation by the Gene ID. Then, for each protein-coding gene in GENCODEv19, we extract its transcription start site (TSS) and extend it by 1.5KB (default value, modifiable by the user) upstream and downstream to include potential alternative transcription start sites of protein-coding genes. Any putative lncRNA overlapping with these regions on the same strand will be excluded from the putative lncRNA list considering that alternative TSSs of the protein-coding genes may appear as novel transcripts. Annotated microRNAs, snRNAs, srpRNAs, tRNAs, scRNAs and antigen receptors datasets were obtained from UCSC and ENSEMBL databases. The last step of filtration is to exclude transcripts less than 200nt (default value, modifiable by the use) in length, based on the lncRNA definition. This subset of putative lncRNA is considered to be the comprehensive *putative* lncRNAs assembly and is used in all downstream analyses. Moreover, we annotate all the remaining putative lncRNAs into different categories based on their overlaps with RefSeq, ENSEMBL and MiTranscriptome.

### Estimation of lncRNAs abundances

We use featureCounts (27) to count the raw reads for all the genes included in the putative lncRNA assembly and GENCODEv19 annotations. Strand-specificity and counting single-end versus paired-end reads is determined by the pipeline automatically in the previous step. The read abundance calculation is performed at the gene level and all the reads included are uniquely mapped. FPKM values are calculated accordingly. A summary bar chart is automatically generated after featureCounts is performed (example in Error! Reference source not found.), showing the number and percentage of reads/fragments that have been utilized by the assembly. Problematic samples can be identified in this step and should be excluded from the study if the percentage of reads/fragments assigned to the assembly is too low.

### Coding potential estimation

We use Coding Potential Assessment Tool (CPAT) to estimate the coding potential of the filtered putative lncRNA. Using the pre-trained human (hg19) logistic regression model, CPAT reports putative ORF size and coding probability for each transcript. The optimum cutoff for human gene annotation is 0.364 as determined in the CPAT manuscript. The distribution of ORF sizes and coding potential scores for protein-coding and non-coding transcripts is produced automatically (example shown in Figure 4; see results for details).

**Figure 4.**
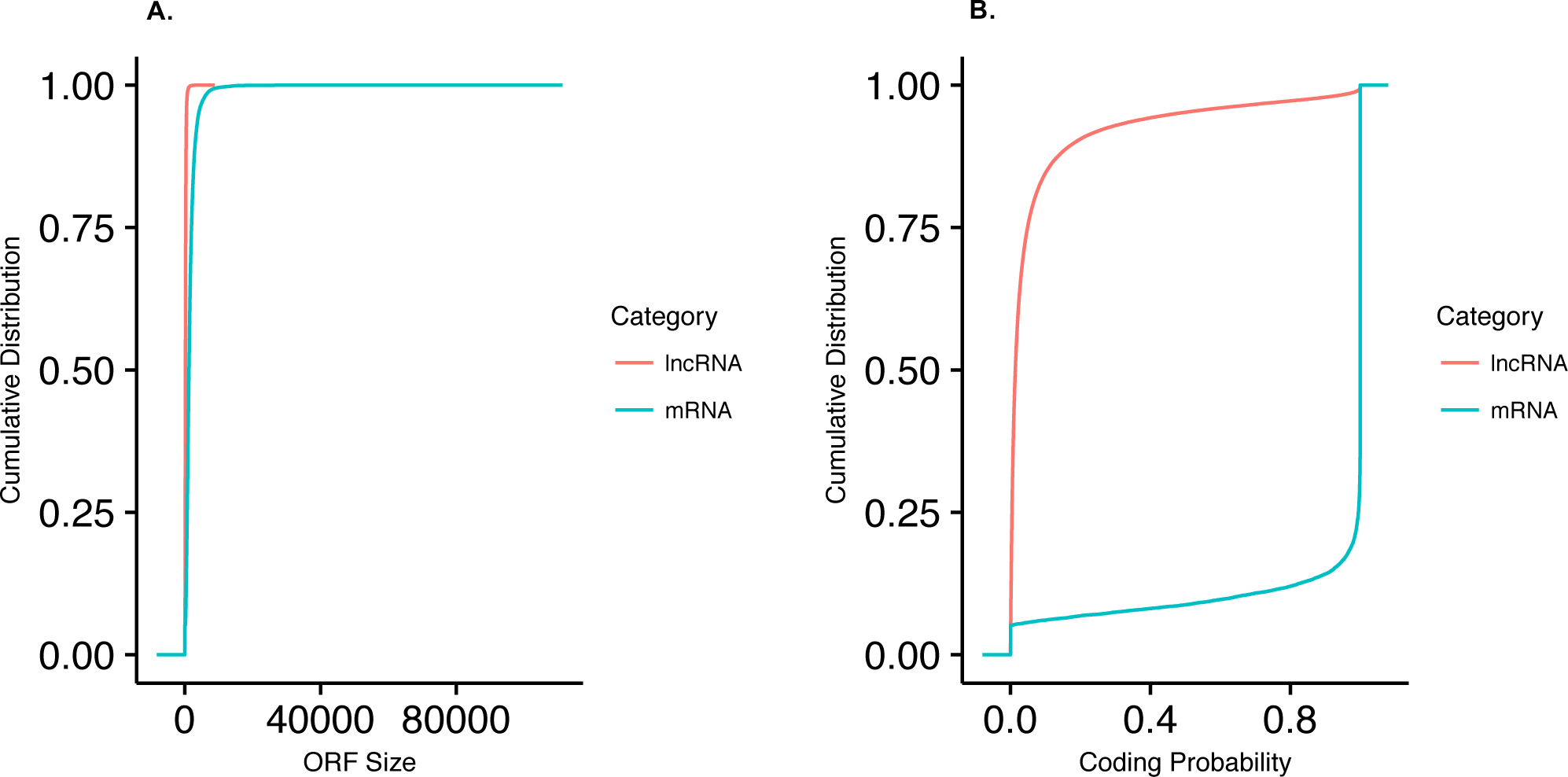
Coding Potential Statistics. (A) Distribution of ORF size for novel lncRNAs, annotated lncRNAs and protein coding genes. (B) Distribution of coding potential score calculated by CPAT for novel lncRNAs, annotated lncRNA and protein coding genes.

### Integration of histone mark ChIP-seq data

ChIP-seq peaks for H3K4me3, H3K27ac and H3K4me1 can be used directly as input to lncRNA-screen. Alternatively, lncRNA-screen can perform its own ChIP-seq analysis starting from raw fastq files. Using Botwie2 (28) for alignment and Macs2 (29) for peak calling, histone mark peaks can be identified. Broad peak calling is used (q-value < 0.05) and fold enrichment compared to the input is calculated. A user-defined fold change cutoff can be applied. Then we extend each putative lncRNA transcription start site by 2.5KB upstream and downstream and assign to it the histone marks that have a peak overlapping this extended TSS region. The fold enrichment value of each overlapping peak is reported for each lncRNA.

### Defining group-enriched lncRNAs

In order to determine all expressed lncRNAs in a specific group, the pipeline allows the users to set a FPKM cutoff based on the distribution of the lncRNA expression level. Then, we compute the average FPKM values for all the putative lncRNAs among different sample groups. Samples having FPKM value below the cutoff are excluded before computing average FPKM value within groups. The number of samples that have an FPKM value above the cutoff will also be reported. Next, combining the ChIP-seq histone mark overlaps with the FPKM cutoff, we are able to define group-enriched expressed lncRNAs based on their expression value and the histone mark enrichment in the extended TSS regions. Users can specify which histone mark (or combination of histone marks) must be present. By default, we define a lncRNA as being expressed in a specific group by requiring the mean FPKM in this group to be greater than a user-defined cutoff (default 0.5) while overlapping with H3K4me3 histone marks in its extended TSS region. Additionally, we define an enhancer-RNA to be expressed in a specific group, if its mean FPKM in this group exceeds the cutoff value while overlapping with both H3K4me1 and H3K27ac but not with H3K4me3 histone marks in its extended TSS region. A “pie matrix” illustrating the number of lncRNAs in categories characterized by different histone marks, as well as the intersections between different sample groups is automatically generated (example shown in Figure 5). The group information table, as described in a previous section, is essential for the pipeline to group the related samples or replicates under the same group name and to match the RNA-seq data and ChIP-seq data. Different levels of grouping can be achieved by adding an additional group column so that users can explore the similarity and difference between samples in different ways.

**Figure 5.**
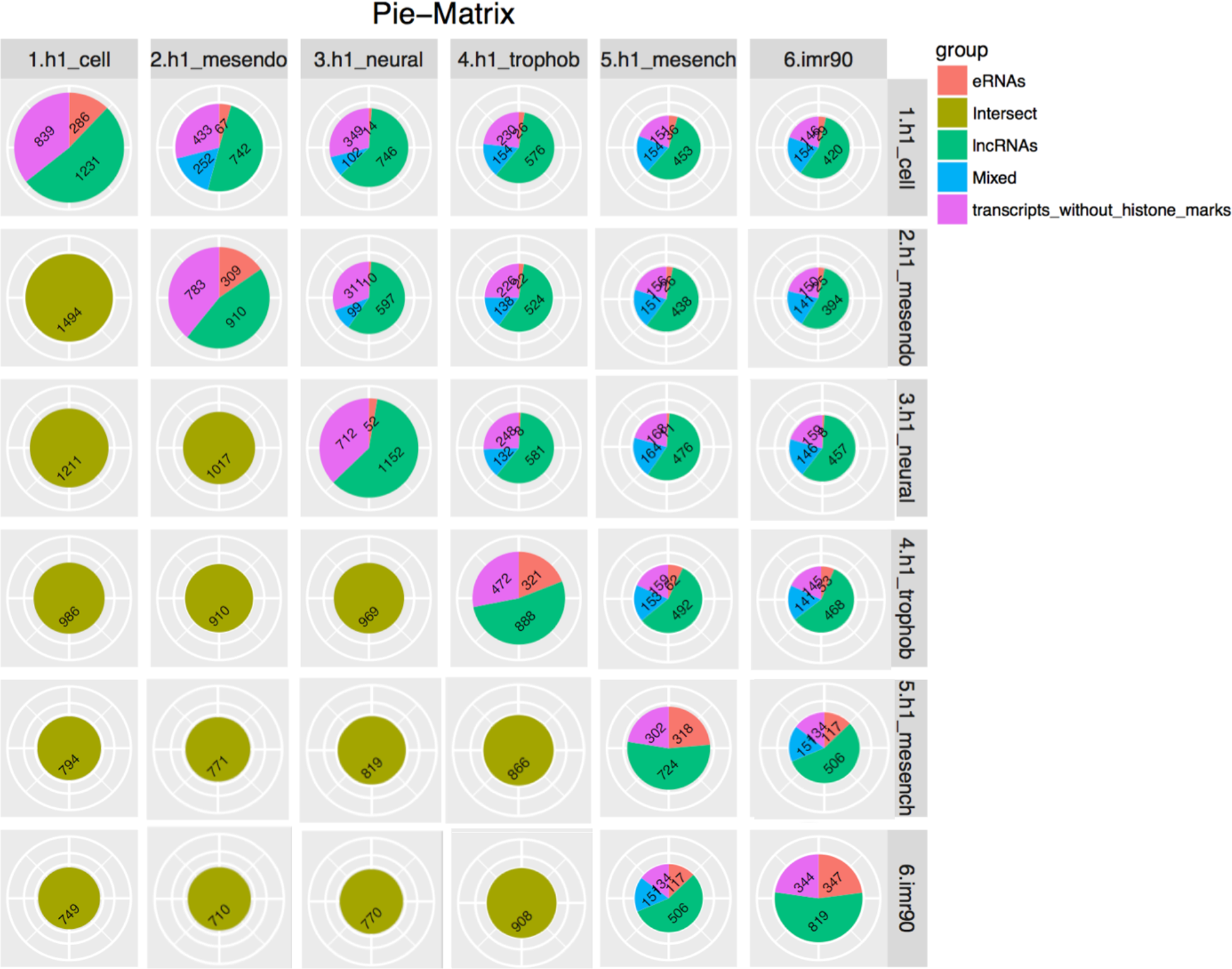
Pie-matrix showing the number of lncRNAs in categories defined by different histone mark enrichment profiles and pairwise overlap of lncRNAs between different groups in different categories. The pie charts in the diagonal from top left to bottom right show total number of lncRNAs in each groups and categorize based on the histone mark enrichment profile. Other pie charts above this diagonal show the overlap of lncRNAs between the column and row names of the groups in different histone mark enrichment categories. Remaining pie charts below the diagonal show only the total number of overlapping lncRNAs between the column and row names of the groups. The sizes of the pie charts represent the total number of lncRNAs in each group.

### Pairwise differential expression analysis between designated groups

Our pipeline also integrates pairwise differential expression analysis using DESeq2 (30). If a list of pairwise comparison groups of interest is provided, the pipeline performs all the differential expression analyses provided in the list. Otherwise, by default the program performs differential expression for up to 10 comparison groups. P-values, FDR, and log2 fold changes are provided for each lncRNA.

### lncRNA selection and the comprehensive putative lncRNA feature report

The default criteria of lncRNA selection are illustrated in the example flowchart of Figure 6. The flowchart is generated automatically and shows in detail the filtering criteria applied in each step as well as the number of putative lncRNA selected (and excluded). This allows the users to inspect the breakdown of the impact of each filtration step. The users can then modify the parameter file and re-run all the steps after the “expression estimation” step, thus obtaining an updated lncRNA list and the corresponding flowchart using their custom criteria. Finally, we collect all the results generated by our pipeline into a comprehensive putative lncRNA feature report which includes the columns shown in Table 3. The comprehensive putative lncRNA feature report is provided to the users in both HTML (example snapshot shown in Figure 9) and Excel formats. Both versions allow users to conveniently filter and search based on user-defined criteria.

**Figure 6.**
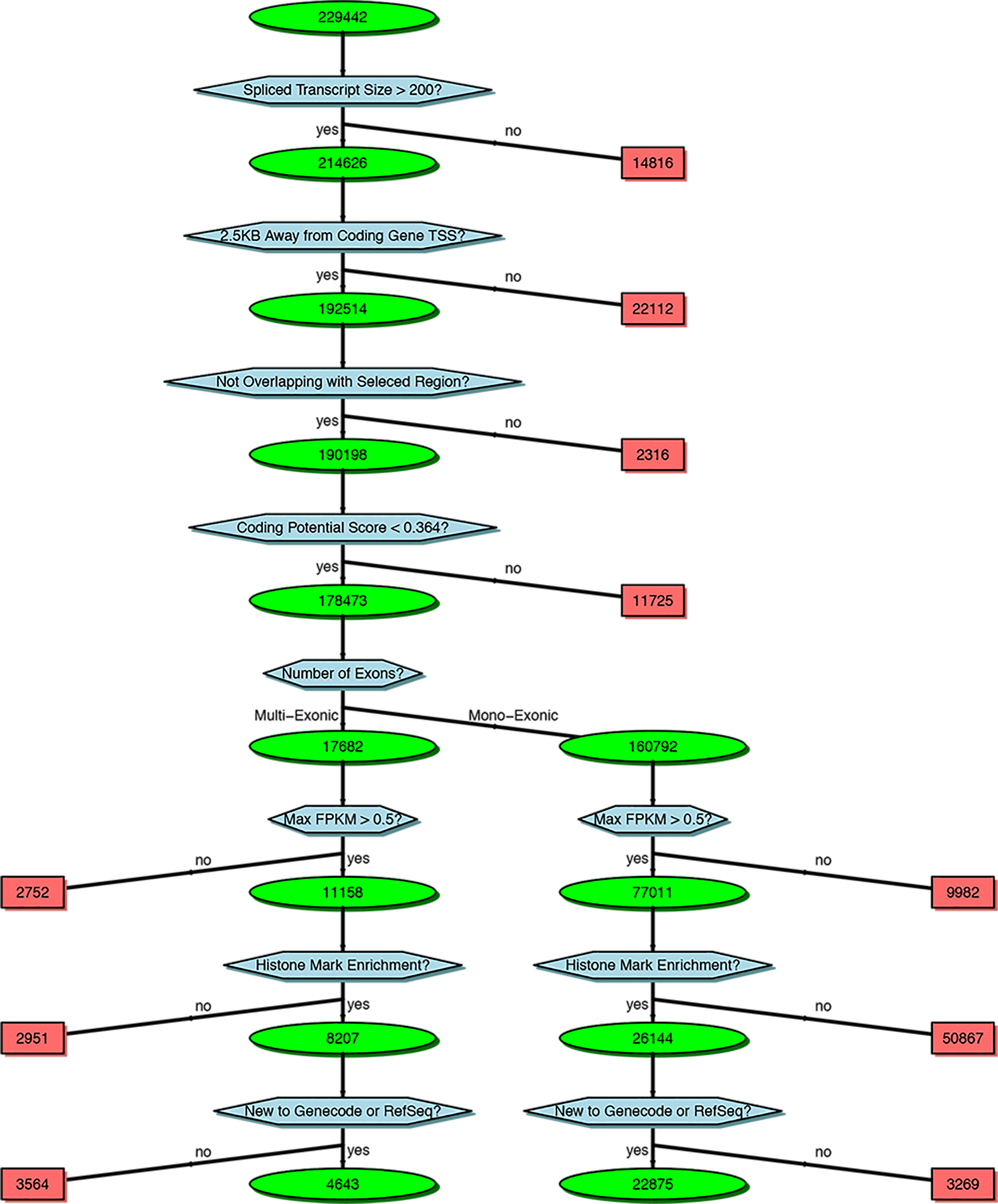
Flowchart of lncRNA selection criteria and number of lncRNAs retained in each step.

### lncRNA heatmaps

Two types of heatmaps are generated for selected lncRNAs which pass all the filtration criteria. First, we generated a supervised clustering heatmap (example shown in Figure 7), designed to show the group-enriched lncRNAs. Because a lncRNA may be expressed in different groups at the same time, in this type of heatmap, the lncRNAs may appear multiple times. The order of the sample groups is the same as the order of the lncRNAs discovered in each group, which ensures that the group-enriched lncRNAs are always located near the diagonal of the heatmap. Second, we provide an unsupervised hierarchical clustering heatmap (example shown in Figure 8) for samples (columns) as well as lncRNAs (rows). This heatmap lets the users inspect the similarity and specificity between sample groups in the filtered lncRNA level, and also allows the users to search for lncRNAs co-expressed or co-differentially expressed between groups.

**Figure 7.**
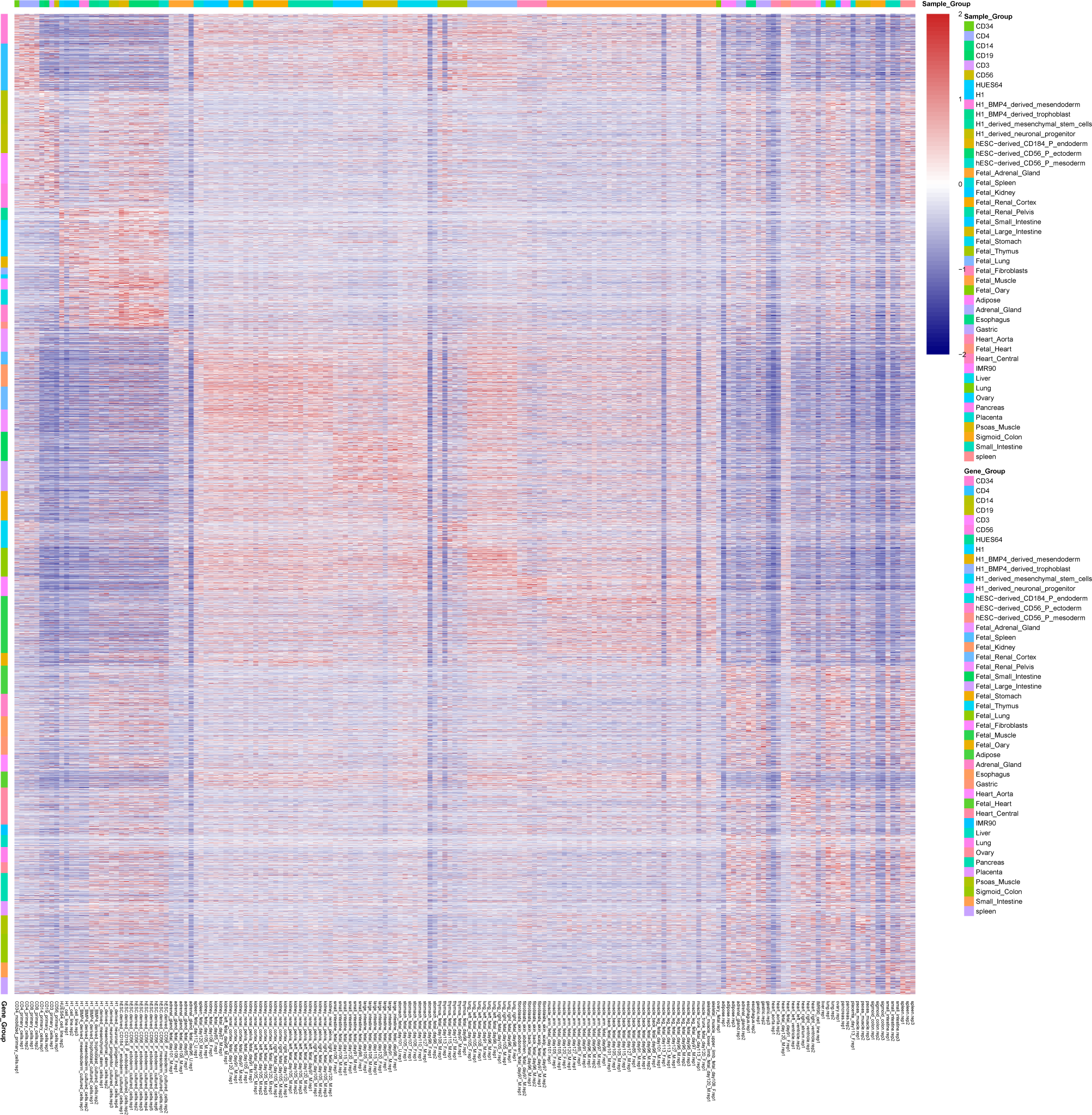
Supervised lncRNA expression heatmap. It plots all group-enriched lncRNAs selected by user-defined filtration criteria. lncRNAs may appear multiple times due to co-expression in multiple groups. The order of the samples is the same as the order of the lncRNA discovered in each group.

**Figure 8.**
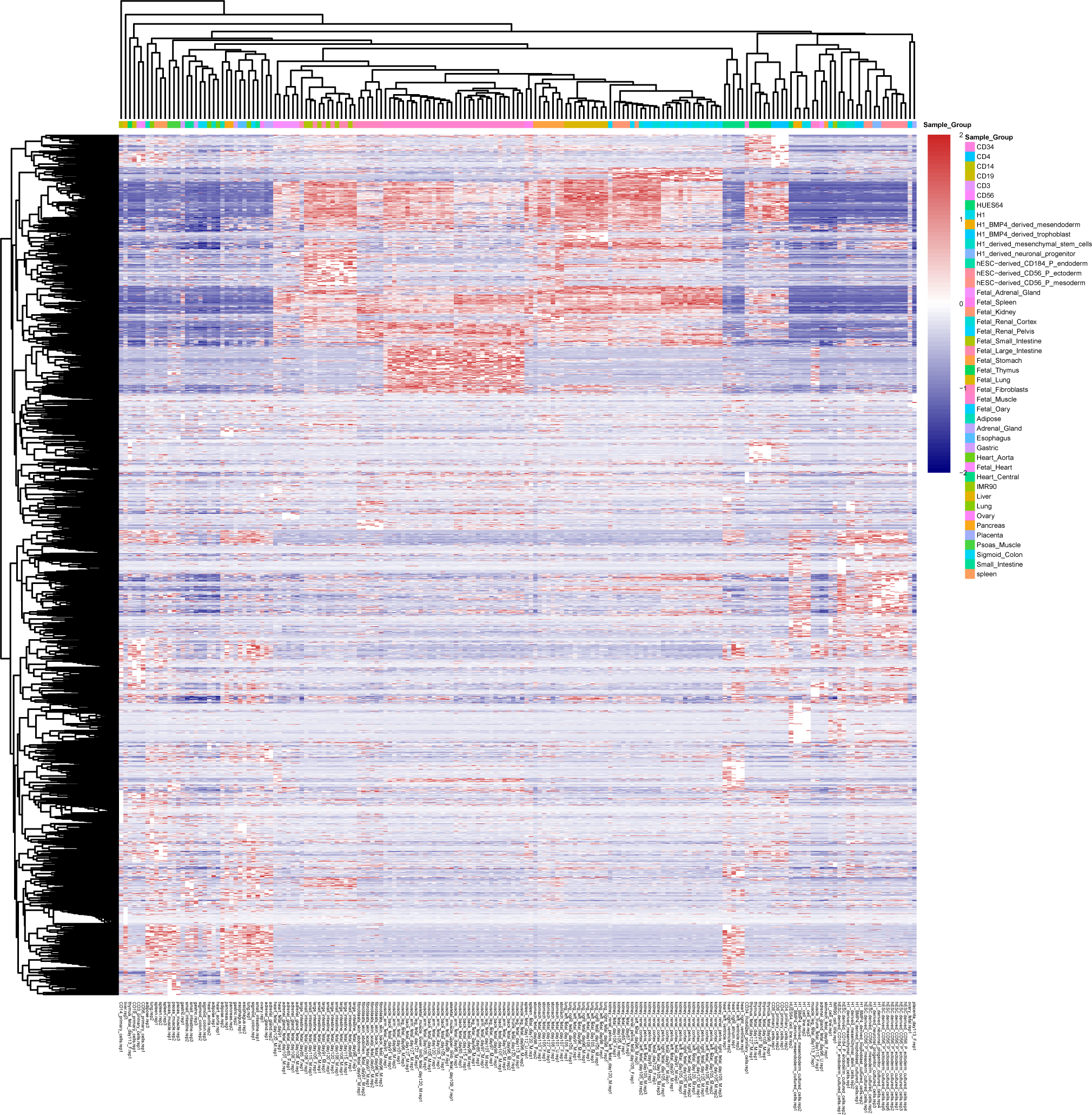
Unsupervised lncRNA expression heatmap. Hierarchical clustering is performed in both rows and columns which categorized lncRNAs into different groups and identify sample similarity and difference.

### lncRNA genome snapshots

The pipeline generates a genome snapshot (example shown in Figure 10) for each lncRNA, centered around the lncRNA locus and zooming out 10 times from the lncRNA length. In this snapshot, users can choose to display ENSEMBL, RefSeq, GENCODEv19, or other user-defined GTF or BED format annotations. The comprehensive merged assembly is also shown in this snapshot, containing all the transcripts assembled without any filtration criteria steps. Moreover, the users can choose to plot bigwig RNA-seq signal tracks either at the merged group level or at the sample level. The HTML report (Figure 9) also includes a preview of the lncRNA genome snapshot for all lncRNAs.

**Figure 9.**
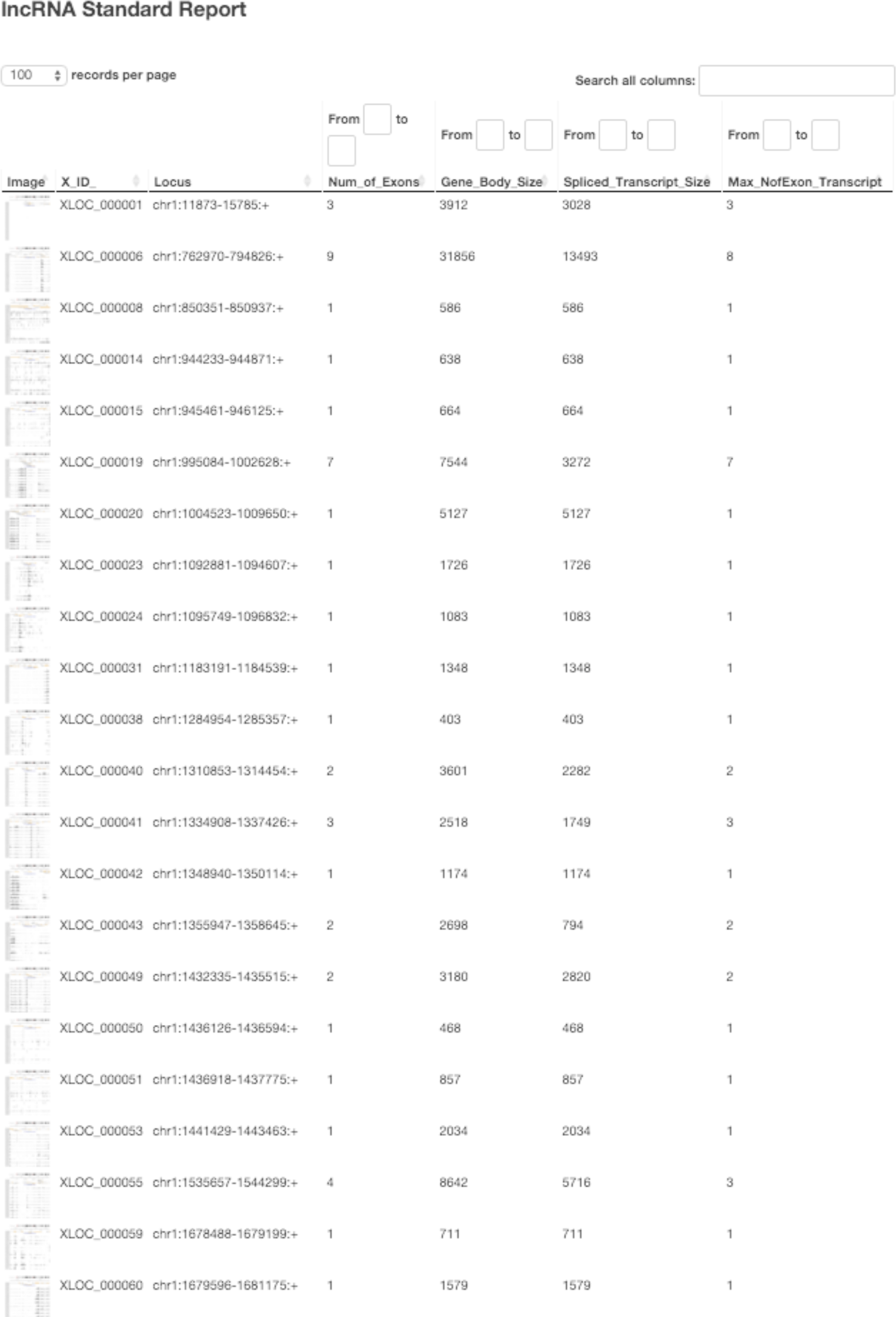
A screenshot of the interactive HTML report generated by lncRNA-screen, which enables sorting, filtering, searching based on different lncRNA features and a preview of the lncRNA genome snapshot and local heatmap.

**Figure 10.**
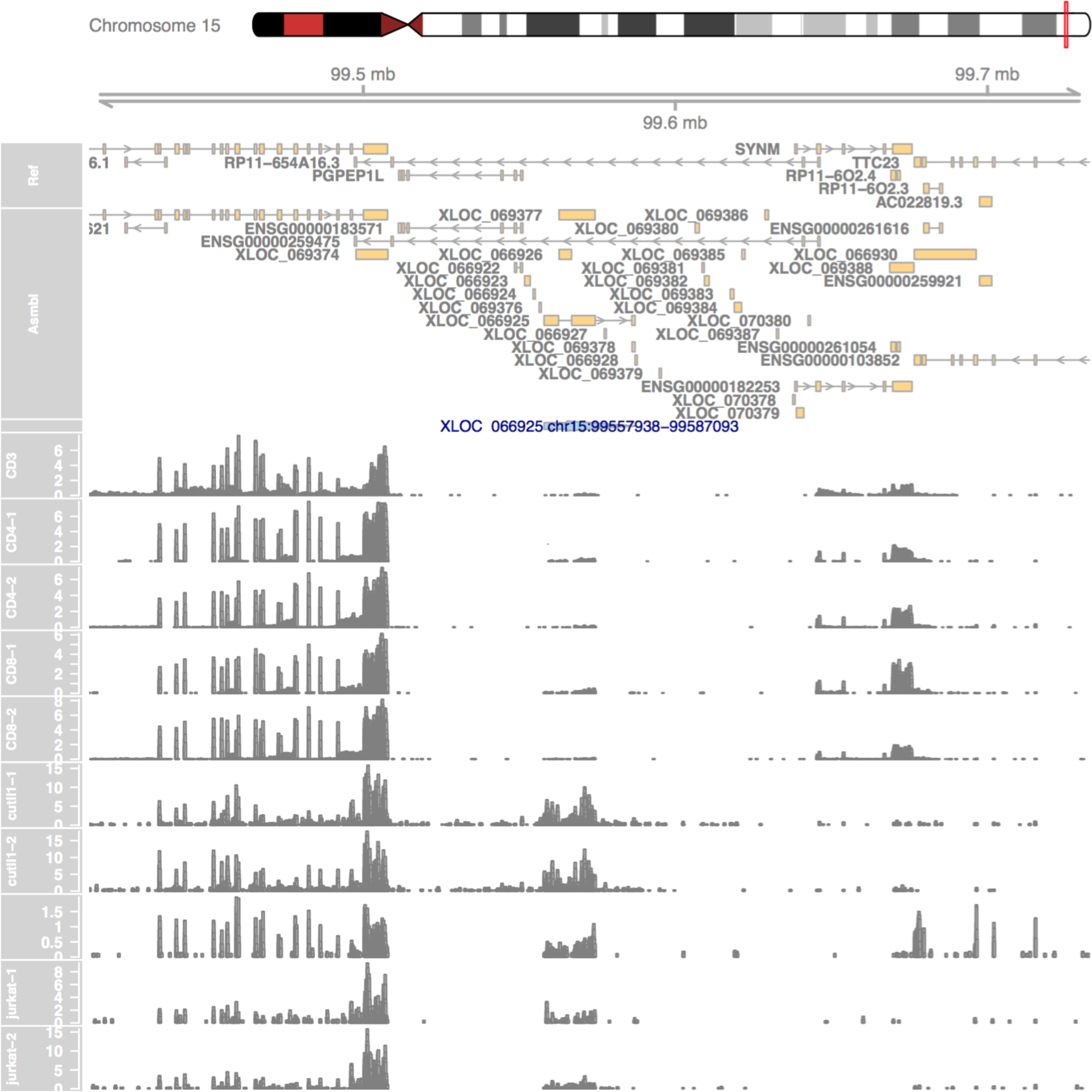
lncRNA genome snapshot. Snapshot of the area surrounding each lncRNA, showing the RefSeq annotations, lncRNA-screen assembly and selected RNA-seq or ChIP-seq signal tracks.

## RESULTS

### Quick installation, setup, execution and interactive browsing of the results

lncRNA-screen can be downloaded as a single zip file from GitHub through the link below. The reference files setup script step will automatically download the necessary references and dependencies. Following instructions in the “how to run” link below, user can easily set up the preferred parameters and run the entire pipeline using a single command. After the run is completed, the user can interactively browse the results using the automatically generated HTML report (see example below) or import the automatically generated table into Excel to do more complex filtering.

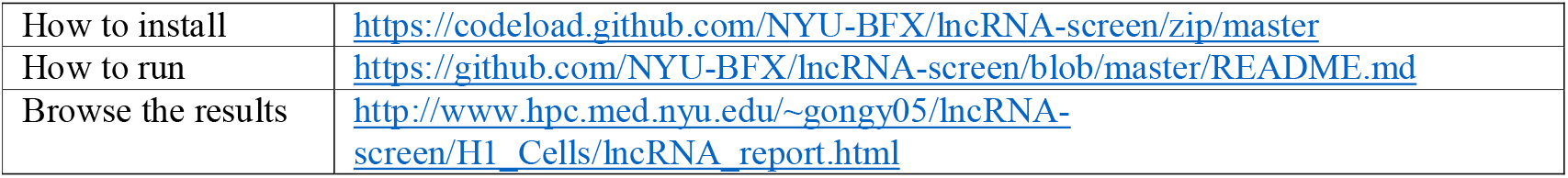

### lncRNAs in Roadmap Epigenomics

To demonstrate usage and performance of lncRNA-screen in big datasets, we used data from the Roadmap Epigenomics project (31), which contains RNA-seq and ChIP-seq data representing a collection of human stem cells and tissue type. The raw FastQ reads of total 198 RNA-seq samples from Roadmap Epigenomics project were downloaded from SRA using SRA-toolkit and aligned to the GENCODEv19 reference genome by STAR (version 2.4.2a) with default parameters (see methods). After quality control (Figure 2 and Error! Reference source not found.), 187 samples (see Error! Reference source not found.) were successfully processed and classified into 40 groups based on cell type. Accepted reads for each sample were assembled individually using Cufflinks (version 2.2.1) providing the guide reference (RefSeq Flat Table GTF file). Cuffmerge was employed to merge all the assemblies into a comprehensive transcriptome assembly, yielding 491218 transcripts, forming 229442 genes in total. By comparing the merged comprehensive transcriptome assembly with GENCODEv19 annotated genes using Cuffcompare, transcripts are classified into different categories based on their structure compatibility of the GENCODEv19 reference annotation. All filtering steps in Phase I of lncRNA-screen, including the number of lncRNAs selected and discarded by various filtering criteria are shown in Figure 6. The total number of putative lncRNAs identified are 178473. Both our coding genes, annotated and novel lncRNA candidates were tested by CPAT and the coding potential distribution comparison are in Figure 4. Novel lncRNA and annotated lncRNA discovered in this pipeline showed similar distribution in ORF and coding potential, and significantly different than coding genes. We use the recommended coding potential cutoff 0.364 for human genome, excluding 11725 genes from our putative lncRNA list, which is 6% of our total candidates. We also included the ChIP-seq histone marks broad peak calling result from macs2 (H3K4me3, H3K27ac and H3K4me1) from Roadmap Epigenomics project for all of the 40 groups and matched them with corresponding RNA-seq sample groups. The lncRNA feature report in both HTML (see the link in github) and Excel format (see the link in github) included all 178,473 putative lncRNAs and their features shown in Table 3. The lncRNA feature report (Figure 9) not only enables sorting, filtering, searching for all lncRNA features extracted from different data resources, but also includes a preview of the lncRNA genome snapshot. Examples of high confident lncRNA local snapshot is shown in Figure 10. The report also includes a pre-set UCSC Genome browser session link to provide advanced genome browse function. With default filtration parameters, pie-matrix (Figure 5) is showing all the lncRNAs expressed (mean FPKM > 0.5) in at least one group by categories defined by 3 histone mark overlaps in the extended TSS regions (TSS flanked by +/- 1.5kb). It also shows pairwise overlap of lncRNAs between different groups in different categories. A total of 8,207 unique lncRNAs were identified as expressed in at least one group and have all H3K4me3, H3K4me1 and H3K27ac histone marks enrichment in its matched group. And total of 26,144 unique enhancer-RNAs were identified as expressed in at least one group and having both H3K4me1 and H3K27ac but don’t have H3K4me3 histone marks enrichment in the matched group. A breakdown list of number of lncRNAs and enhancer-RNAs identified in each group is in Table 2. For these lncRNAs identified in each group, we generated a supervised heatmap (Figure 7) ensuring the order of the groups and the order of lncRNAs of each groups to be identical for rows and columns. Therefore, the diagonal position of the heatmap shows relatively higher FPKM values across all the samples which proved that lncRNAs identified in a specific group have the relatively higher expression values comparing to other groups. And the unsupervised automatic hierarchical clustered heatmap (Figure 8) for all the 8,207 unique group-enriched lncRNAs revealed some clusters of lncRNAs which are differentially expressed in TALL cell lines comparing with other cell types.

**Table 2.** Number of putative lncRNAs and enhancer-RNAs identified in each group.

| Group | Number of lncRNAs |  | Group | Number of lncRNAs |
| --- | --- | --- | --- | --- |
| Adipose | 8709 |  | Fetal Thymus | 5763 |
| Adrenal Gland | 3950 |  | Gastric | 3978 |
| Bladder | 695 |  | H1 BMP4 derived mesendoderm | 1711 |
| CD14 | 3982 |  | H1 BMP4 derived trophoblast | 844 |
| CD19 | 6188 |  | H1 derived mesenchymal stem cells | 443 |
| CD34 | 3661 |  | H1 derived neuronal progenitor | 1067 |
| CD3 | 6113 |  | H1 | 8271 |
| CD4 | 5232 |  | Heart Aorta | 17092 |
| CD56 | 9596 |  | Heart Central | 6126 |
| Esophagus | 3443 |  | hESC-derived CD184 P endoderm | 1367 |
| Fetal Adrenal Gland | 12008 |  | hESC-derived CD56 P ectoderm | 2086 |
| Fetal Fibroblasts | 1568 |  | hESC-derived CD56 P mesoderm | 900 |
| Fetal Heart | 2081 |  | HUES64 | 1257 |
| Fetal Kidney | 4429 |  | IMR90 | 2152 |
| Fetal Large Intestine | 5824 |  | Liver | 785 |
| Fetal Lung | 4930 |  | Lung | 2359 |
| Fetal Muscle | 10477 |  | Ovary | 1056 |
| Fetal Oary | 2472 |  | Pancreas | 5204 |
| Fetal Renal Cortex | 4578 |  | Placenta | 9030 |
| Fetal Renal Pelvis | 4621 |  | Psoas Muscle | 2223 |
| Fetal Small Intestine | 4508 |  | Sigmoid Colon | 18827 |
| Fetal Spinal Cord | 8113 |  | Small Intestine | 4760 |
| Fetal Spleen | 2448 |  | spleen | 2953 |
| Fetal Stomach | 10330 |  | Testes Pool | 1266 |

**Table 3.** List of features included in the final lncRNA feature report.

| Single Column | One column per group |
| --- | --- |
| lncRNA ID | Mean FPKM above user-defined threshold |
| Locus | Percentage of samples above FPKM threshold |
| Number of Exons | H3K27ac peak enrichment score |
| Gene Body Size | H3K4me3 peak enrichment score |
| ORF Size | H3K4me1 peak enrichment score |
| Coding Probability | Histone marks enrichment and FPKM cutoff combination |
| Gencode Annotation | Differential expression analysis between groups |
| RefSeq Annotation | Hi-C interaction between lncRNA and neighboring coding genes |
| Ensembl Annotation |  |
| MiTranscriptome Annotation |  |
| SNPs annotation |  |
| Copy number gain/loss value |  |

### Integration with Hi-C data

To demonstrate the flexibility of our pipeline we integrated RNA-seq/ChIP-seq data from Roadmap Epigenomics with matched Hi-C data from this study (32, 33). First, we rerun the lncRNA-screen pipeline focusing only on the samples that have matched Hi-C data (the report is included in github). For each cell type, we defined expressed mRNAs, lncRNAs including annotated and novel, enhancer-RNAs and enhancer regions using expression profile and H3K4me3, H3K27ac and H3K4me1 histone mark occupancy. The numbers of elements in each category for each cell type are included in Table 4. Hi-C analysis was performed using our HiC-bench pipeline (34). HiC-bench automatically produces various plots to help the user assess the quality of the data as well as compare different samples. Paired-end reads were mapped to the reference genome (hg19 or mm10) using Bowtie2 (28). Local alignments of input read pairs were performed as they consist of chimeric reads between two (non-consecutive) interacting fragments. This approach yielded a high percentage of mappable reads (>25%) for all datasets. Mapped read pairs were subsequently filtered for known artifacts of the Hi-C protocol such as self-ligation, mapping too far from the enzyme’s known cutting sites etc (Figure 11). Samples clustered as expected by Principal Component Analysis (Error! Reference source not found.**A**), and the average Hi-C count showed the characteristic dependency on the distance between the interacting fragments as demonstrated in previous studies (Error! Reference source not found.**B**). **Error! Reference source not found.A,B** shows the sizes and the number of detected topologically-associated domains (TADs) detected in each cell type and each replicate, and Error! Reference source not found.**C** shows the pairwise overlaps of boundaries between all pairs of samples and replicates. HiC-bench also generates a table of all the interacting loci annotated with genes, ChIP-seq peaks and any other region file that the user provides. Using this feature, we compiled a comprehensive report of all interactions that involve the lncRNAs discovered by our lncRNA-screen pipeline in the matched RNA-seq/ChIP-seq datasets. Overall, we found that 268 lncRNAs in hESC, 239 lncRNAs in mesendoderm cells, 9 lncRNAs in neural progenitor cells and 254 lncRNAs in mesenchymal cells interacting with at least one mRNAs in cis within the context of topological domains. Most importantly, mRNA expression appears to be sensitive to changes in looping with their lncRNA interacting partners. We used HiCPlotter (35) to generate the Hi-C maps mRNA-lncRNA interaction plots. As an example we show putative lncRNA CTD-2006C1.2 on chromosome 19 in Figure 12. This lncRNA interacts with multiple protein-coding genes, mainly zinc-finger proteins within a single topological domain. When we compared both the expression and the Hi-C interaction intensity of the lncRNA to its neighboring genes, we observed an interesting contrasting pattern. Upstream, the lncRNA interacts with two protein-coding genes, ZNF441 and ZNF491: both genes are upregulated in Mesenchymal cells compared to hESCs and show a concomitant increase in Hi-C looping. In contrast, downstream, the lncRNA interacts with ZNF878 and ZNF625, both downregulated in Mesenchymal cells compared to hESCs with concomitant decrease in Hi-C looping.

**Table 4.** Number of transcripts by category for each multi-lineage differentiated embryonic stem cells.

| Cell Type | enhancers | eRNA | lncRNA | mRNA |
| --- | --- | --- | --- | --- |
| hESC | 6323 | 286 | 1231 | 8605 |
| Mesench | 16736 | 318 | 724 | 8660 |
| Mesendo | 2371 | 309 | 910 | 8396 |
| Neupro | 1487 | 52 | 1152 | 8652 |
| Trophect | 12663 | 321 | 888 | 8766 |

**Figure 11.**
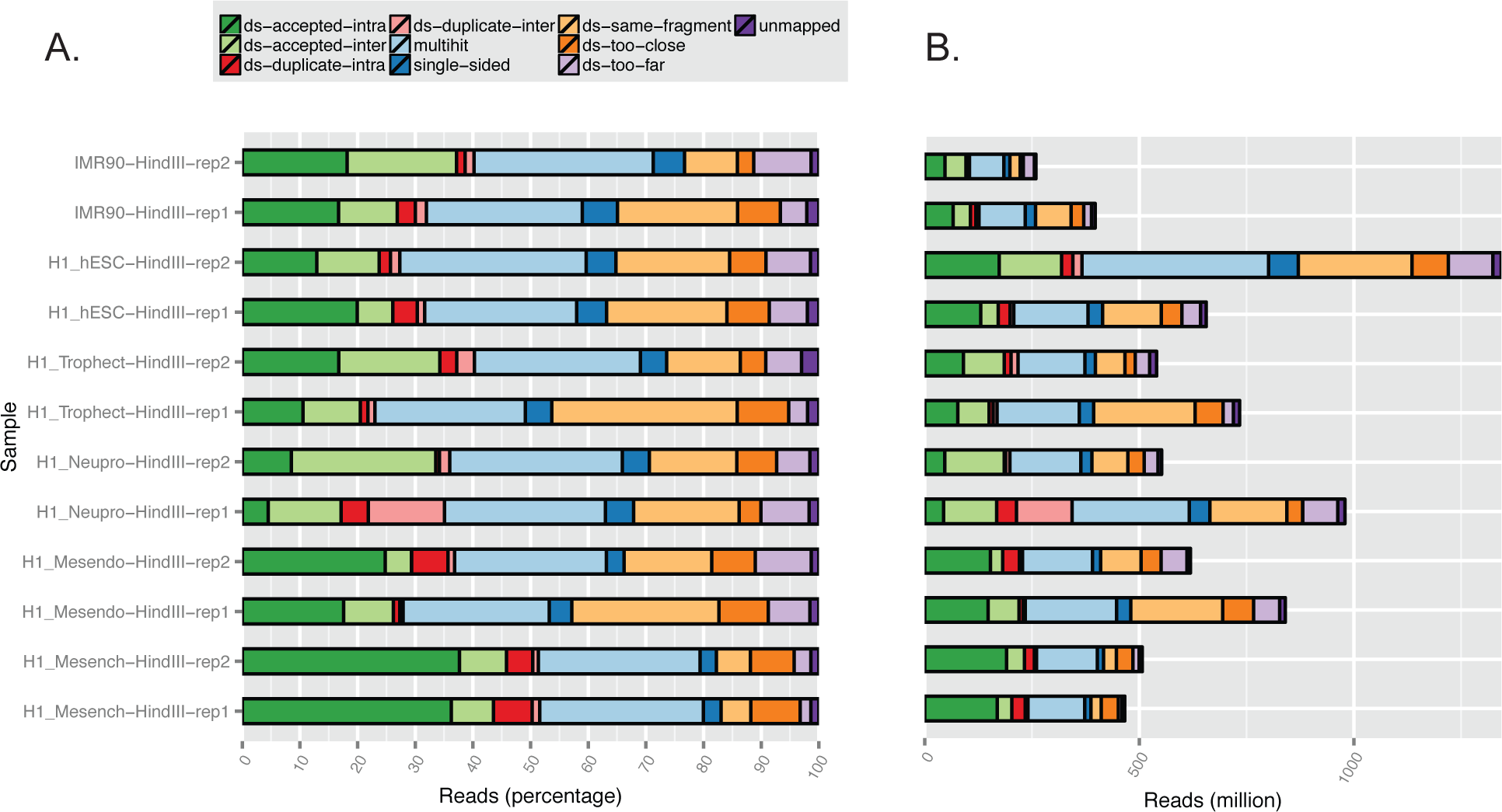
Hi-C quality assessment plots. (A) Alignment and filtering statistics (read pair counts and percentages). (B) Average normalized Hi-C counts are presented as a function of the distance between the interacting partners.

**Figure 12.**
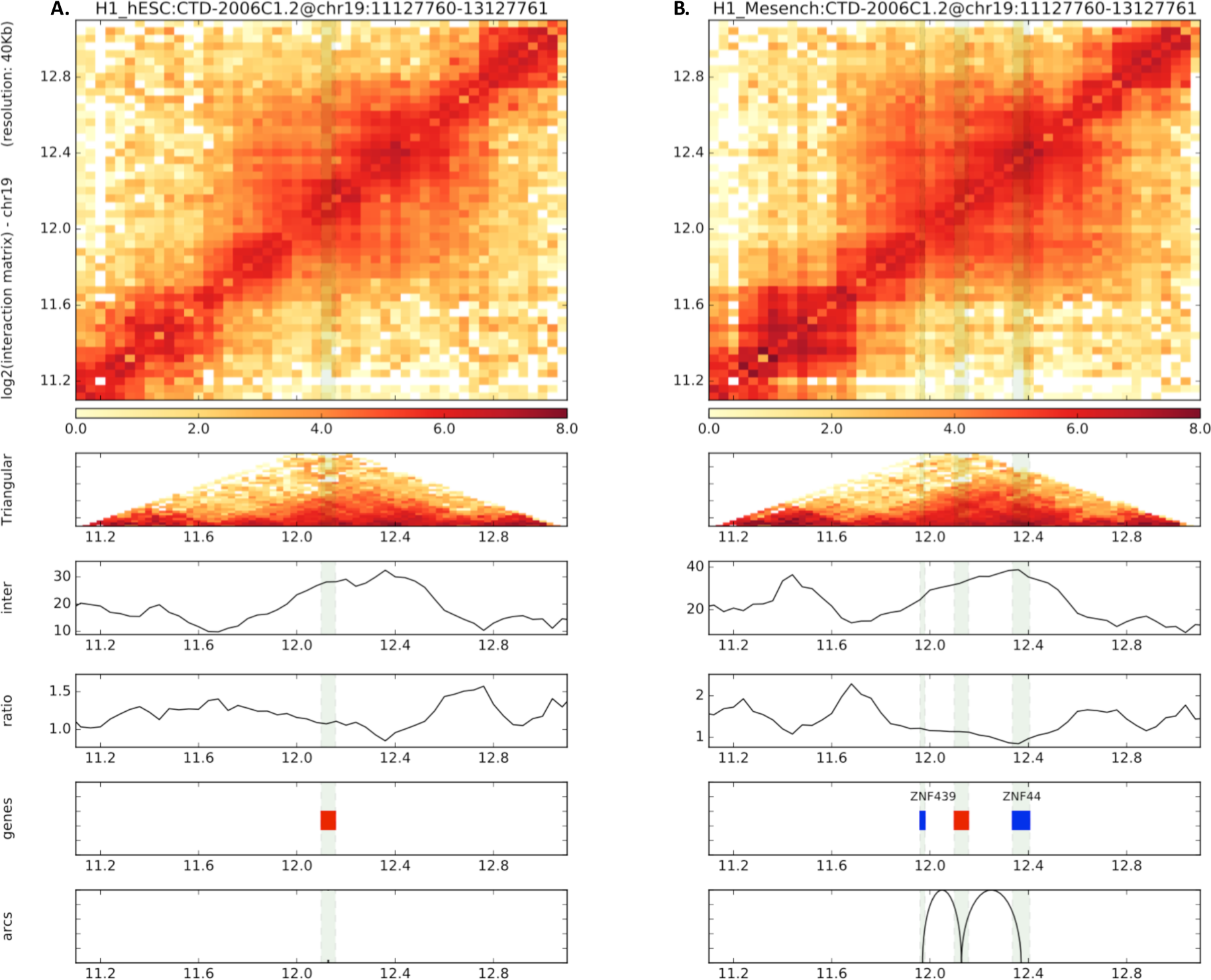
Example of lncRNA-mRNA interactions. Putative lncRNA CTD-2006C1.2 is shown to differentially interact with two upstream and two downstream zinc-finger genes: hESCs (left panel) and Mesenchymal (right panel).

## DISCUSSION

We developed lncRNA-screen, an easy-to-use integrative lncRNA discovery platform for comprehensive mapping and characterization of lncRNAs using a variety of genomics datasets. The main objective of this work was to facilitate the computational discovery of lncRNA candidates to be further examined by experimental screening as well as functional experiments. More specifically, our goal was to enable experimental laboratories with limited genomics expertise to quickly and comprehensively characterize lncRNAs in their particular field of study (e.g. cancer, stem cells, development). Our pipeline can be installed using one self-contained installation package available on GitHub, and, importantly, is designed to *enable execution of an entire analysis using a single command*. Initializing a new analysis is also simplified into setting up a trivial sample sheet to describe the datasets involved in the study. Additionally, lncRNA-screen generates an interactive lncRNA report which allows users to explore the results of the analysis, define their own custom criteria for selecting lncRNAs, and interactively visualize the filtered results using the UCSC Genome Browser and pre-built genome snapshots. The pipeline is compatible with both stand-alone server environments and high-performance computing clusters. Although lncRNA-screen provides a complete and convenient method for lncRNA discovery to the scientific community, there are several challenges and improvements to be addressed in the future. For example, although cufflinks has proved to be an accurate assembler for RNA-seq data (especially protein-coding genes), the lower levels of lncRNA expression may lead to inaccurate or incomplete transcriptome assemblies. In summary, our pipeline provides a comprehensive solution for lncRNA discovery and an intuitive interactive report for identifying promising lncRNA candidates. lncRNA-screen is available as free open-source software on GitHub and our bioinformatics team offers installation and usage support.

## ACKNOWLEDGEMENT

Aristotelis Tsirigos was supported by a Research Scholar Grant, RSG-15-189-01 ‐ RMC from the American Cancer Society. We thank Igor Dolgalev and Stephen Kelly from the NYU Applied Bioinformatics Center and all the Aifantis Lab members for inspiring discussions. We also thank Charalampos Lazaris for advice on how to use HiCPlotter. This work has used computing resources at the High Performance Computing Facility (HPCF) of the Center for Health Informatics and Bioinformatics at the NYU Langone Medical Center, so we thank Eric Peskin, Ali Siavosh-Haghighi and Loren Koenig for their technical support.

## FUNDING

The study was supported by a Research Scholar Grant, RSG-15-189-01 ‐ RMC from the American Cancer Society to Aristotelis Tsirigos (AT) and the reseach program R01CA194923 and R01CA169784 from National Institute of Health (National Cancer Institute) to Iannis Aifantis (IA).

## AUTHOR CONTRIBUTIONS

YG designed and implemented the pipeline, performed computational analyses, generated figures and wrote the user manual. HH and TT offered expertise on lncRNA biology. TT designed an early prototype of the pipeline. AT analyzed the Hi-C data. YG and AT wrote the manuscript. AT and IA designed and supervised this research project.

